# O-GlcNAc Transferase Regulates GABAergic Synapse Organization and Receptor Composition

**DOI:** 10.1101/2025.11.21.689700

**Authors:** Manish Bhattacharjee, Linkun Han, Petya Elenska, Mario Pérez Del Pozo, Carla Blanco, Michael Druzin, Olof Lagerlöf

## Abstract

Neural circuits must integrate metabolic information to maintain stable activity and appropriate behavioral responses. While metabolic regulation of excitatory synapses has been well studied, far less is known about how inhibitory synapses respond to changes in energy state. In this study, we show that O-GlcNAc transferase (OGT), a dynamic sensor of cellular nutrient flux, localizes to postsynaptic sites of the inhibitory synapses where it modulates synapse morphology and receptor composition. OGT over-expression reduced the size and intensity of vGAT and gephyrin puncta, whereas conditional OGT deletion produced a converse enlargement and redistribution of inhibitory scaffolds and vesicular proteins. Furthermore, OGT deletion accelerated inhibitory postsynaptic current decay kinetics and induced subunit-specific shifts in GABA_A_ receptor surface expression: β3 subunits decreased, whereas γ2 subunit total and surface levels increased. Together, these findings identify OGT as a metabolic regulator that modulates inhibitory synapse structure and signaling, providing a mechanistic link between energy state and GABAergic circuit function in health and disease.

## Introduction

Neural circuits in the brain continually sense fluctuations in the metabolic state of the body to coordinate behavior and physiology [1, 2]. If they fail to integrate nutrient and hormonal fluctuations, it can lead to widespread dysfunction across multiple levels of physiology and behavior. Impaired coupling between metabolic state and neuronal activity contributes not only to metabolic disorders such as obesity, metabolic syndrome, and type-2 diabetes[3, 4], but also to neuropsychiatric, neurodevelopmental and neurodegenerative conditions including anorexia nervosa, autism spectrum and Alzheimer’s disease [5–7]. A major mechanism by which neural circuits integrate metabolic information is synaptic plasticity, allowing neurons to adjust synaptic strength and connectivity in ways that stabilize circuit activity and adapt behavior to internal state[8–10]. Identifying the molecular pathways that connect metabolic signaling to synaptic regulation will reveal how energy state influences circuit function and a wide range of disorders[11–13].

Most studies examining how metabolism influences synaptic function have focused on excitatory synapses[14–17]. While excitatory synapses are targets for some metabolic signals that act within neural circuits, far less is known about how inhibitory synapses integrate metabolic information.

Inhibitory synapses are essential for controlling neuronal excitability, synchronizing network oscillations, and stabilizing circuit output[18]. Mediated primarily by γ-aminobutyric acid (GABA) signaling, inhibitory synaptic activity prevents the pathological hyperexcitability associated with epilepsy, schizophrenia, and autism spectrum disorders[19–22]. Beyond maintaining stability, inhibitory synapses exhibit plasticity: activity-dependent changes in GABA_A_ receptor trafficking and remodeling of scaffold proteins such as gephyrin and collybistin enable dynamic tuning of inhibitory strength. This inhibitory plasticity complements excitatory plasticity and contributes to learning, homeostasis, and adaptive circuit reconfiguration [8, 23, 24]. Despite their importance, the molecular mechanisms through which inhibitory synapses are regulated remain poorly understood. Because inhibitory neurotransmission plays a central role in circuit computation and information flow[25], identifying pathways that link metabolic sensing to inhibitory synaptic remodeling is critical for revealing how neural circuits coordinate behavior and physiology[24, 26].

Here, we identify a metabolic pathway that regulates inhibitory synapse organization and receptor composition. O-GlcNAc transferase (OGT) is an intracellular enzyme that attaches N-acetylglucosamine to serine and threonine residues on target proteins in response to glucose and metabolic hormone fluctuations[27, 28]. Dysregulation of OGT has been linked in humans to several disorders associated with impaired brain function, including diabetes, intellectual disability and neurodegenerative disorders[29–31]. Our data show that OGT localizes to inhibitory postsynaptic sites. Manipulating OGT altered the clustering of synaptic markers and the composition of GABA_A_ receptor subunits, leading to changes in inhibitory current kinetics. Using both gain- and loss-of-function approaches, we find that OGT bidirectionally regulates inhibitory synapse structure without altering synapse number. These findings indicate that OGT has a diverse and widespread effect on inhibitory synapse organization. Linking metabolic signaling to inhibitory synapse regulation, this work reveals a molecular mechanism by which metabolic state regulates inhibitory synapse control in health and disease [19, 21].

## Results

### Deleting OGT increases neuron disinhibition

We cultured primary cortical neurons from OGT-floxed mice to investigate the role of OGT in neuronal function. OGT was conditionally deleted from these cultures using a lentivirus expressing both GFP and Cre, while control cells were transduced with a virus expressing only GFP, as previously described [16]. Wild-type (WT) and OGT knockout (OGT-KO) neurons had no noticeable morphological differences (Figure 1A). The loss of OGT had no significant effect on neuron number (Figure.1B; P = 0.505). In contrast, deletion of OGT markedly decreased c-Fos, an immediate early gene induced by neuronal firing and synaptic input integration[32, 33]. We observed that OGT-KO neurons displayed a 78% reduction in c-Fos protein levels relative to WT controls under basal conditions (Figure 1C, D; P = 0.0001). Our and others’ data have shown previously that deleting OGT decreases excitatory synapse number and the synaptic abundance of the GluA2 and GluA3 subunits of the neurotransmitter receptor conducting the majority of the fast excitatory neurotransmission in the brain, the

**Figure 1.**
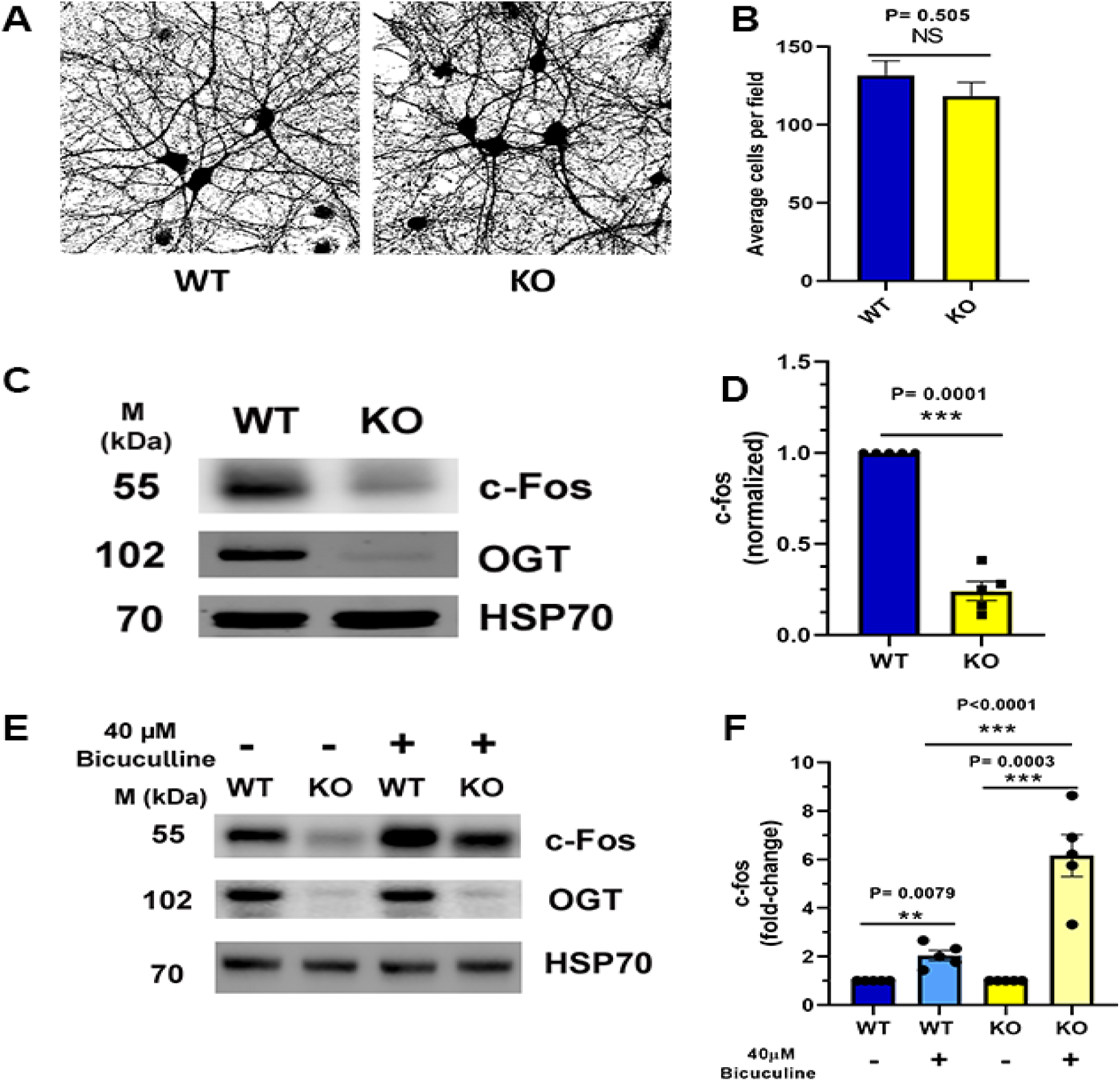
Deleting OGT increases neuron disinhibition. **A**. Thresholded images of GFP expression in WT and OGT-knockout (KO) neurons. **B**. Quantitative analysis of cell viability showing average cells per field in WT and OGT-KO conditions. NS, not significant (P = 0.505) (N= 316(WT) and 301(OGT-KO). **C.** Representative Western blots showing c-Fos, OGT, and HSP70 protein levels under basal conditions. **D.** Quantification of normalized c-Fos expression showing significant reduction in OGT-KO compared to WT neurons under basal conditions (P = 0.0001; N=5). **E**. Representative Western blot analysis of c-Fos protein expression in WT and OGT-KO neurons following treatment with bicuculline (40 μM), with OGT and HSP70 included as controls. **F.** Fold-change analysis of c-Fos expression in WT neurons following bicuculline treatment, with values normalized to WT vehicle control(N=5). Data are presented as mean ± S.E.M (NS, not significant; **P < 0.01; ***P < 0.001, unpaired Student’s t test).

AMPA receptor [16]. Lower excitatory input would decrease c-Fos expression. Alternatively, decreased c-Fos expression could also be an effect of increased inhibitory neurotransmission. We probed inhibitory regulation directly by treating WT and OGT-KO cultures with bicuculline, a GABA_A_ receptor antagonist that blocks inhibitory neurotransmission. Bicuculline increased c-Fos levels in WT neurons as expected (Figure 1E, F). Surprisingly, OGT-KO neurons exhibited a dramatically enhanced response, with c-Fos levels increasing three-fold relative to WT neurons (Figure 1F; P = 0.0003). Together, these findings indicate that OGT deletion produces reduced basal activity-dependent gene expression alongside heightened responsiveness to disinhibition. Our findings support the hypothesis that OGT regulates not only excitatory synapses but also inhibitory synapse function.

### OGT localizes to inhibitory synapses

To investigate whether OGT contributes to inhibitory synaptic regulation, we examined OGT’s subcellular localization in cultured hippocampal neurons. Neurons were transfected at DIV14 with a GFP reporter and immunostained at DIV16 for OGT together with inhibitory synapse markers. The vesicular GABA transporter (vGAT), a presynaptic marker of GABAergic synaptic terminals[34], and gephyrin, the principal postsynaptic scaffold at GABAergic synapses that directly interacts with GABA_A_ receptors [35, 36] were selected as complementary markers of inhibitory synapses.

Confocal microscopy analysis revealed that OGT was prominently detected in dendrites and dendritic spines, consistent with previous reports demonstrating OGT enrichment in the postsynaptic density of excitatory synapses[16]. Importantly, OGT immunoreactivity in dendrites also showed colocalization with both gephyrin-positive postsynaptic clusters (Figure 2A) and vGAT-positive presynaptic boutons apposed to dendrites (Figure 2B). These findings demonstrate that OGT is localized in the postsynaptic terminal of inhibitory synapses, consistent with a role in modulating inhibitory transmission [37].

**Figure 2.**
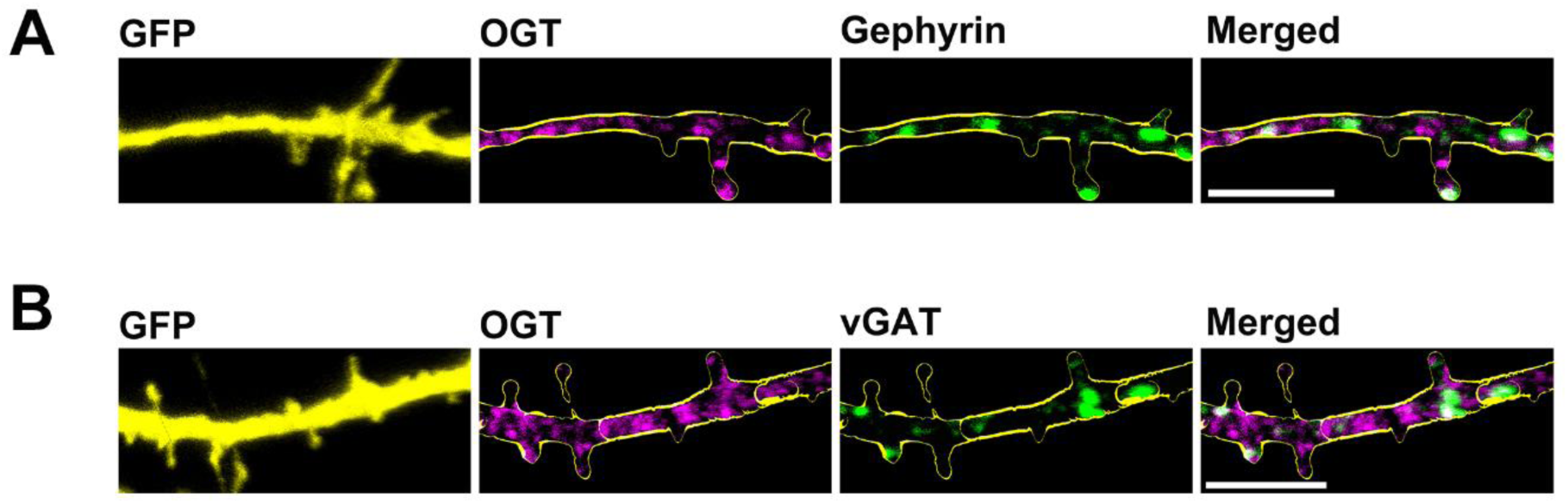
OGT localizes to inhibitory synapses. **A**, Immunostaining of the subcellular localization of OGT (lilac) and gephyrin (green) in cultured cortical neurons (transfected at DIV14). **B**, Immunostaining of the subcellular localization of OGT (lilac) and vGAT (green) in cultured cortical neurons (transfected at DIV14). GFP(yellow) stained the overall morphology of neurons. The scale bars represent 5 μm magnification image.

### OGT overexpression reduces inhibitory synapse size

To systematically test OGT’s role at inhibitory synapses, we first investigated the effects of OGT overexpression on inhibitory synapse morphology and number by transfecting GFP and OGT in cultured hippocampal neurons and immunostaining cells with antibodies against the inhibitory presynaptic marker vGAT and the postsynaptic marker gephyrin. Upon overexpressing OGT in sparse neurons, inhibitory synapses terminating on the overexpressed neuron were identified using 3D Imaris reconstruction software by measuring vGAT and gephyrin puncta intensity, size and number. To illustrate how these inhibitory synapse parameters were defined, Figure 3A shows a representative 3D-rendered dendritic segment depicting colocalized, non-colocalized, and overlapped vGAT and gephyrin puncta used for quantitative analysis.

**Figure 3.**
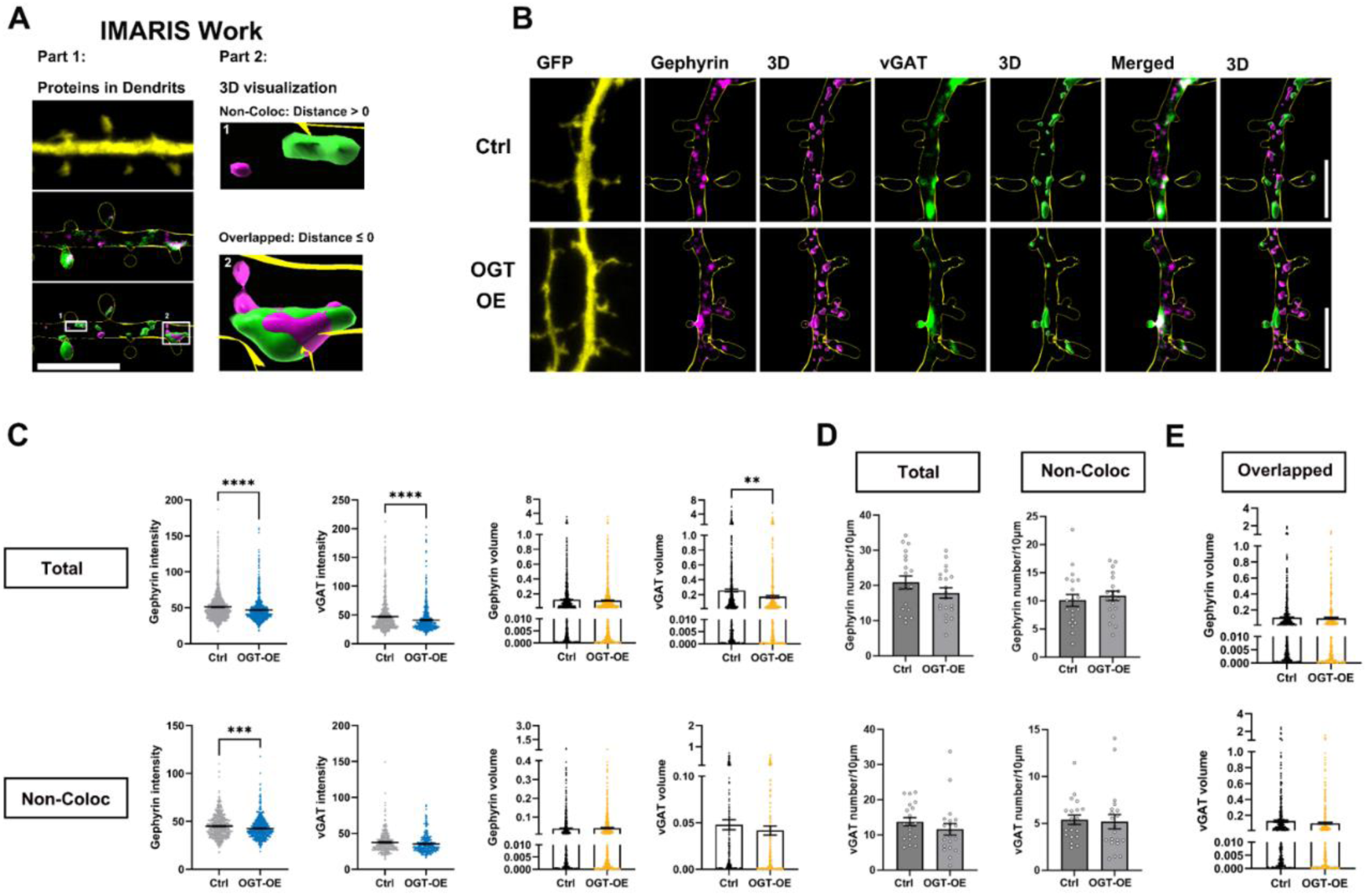
The effect of OGT overexpression on inhibitory synapses. **A**. Representative 3D-rendered dendritic segment illustrating how vGAT and gephyrin puncta were categorized for analysis, including colocalized, non-colocalized, and overlapped populations. **B.** Representative images of cultured hippocampal neurons transfected at DIV7 with expressing GFP alone (Control) or co-expressing GFP with OGT (OGT overexpression). Neurons were analyzed by triple immunofluorescence labeling for GFP (yellow), gephyrin (lilac), and vGAT (green). Scale bar, 5 μm (applies to all images). **C.** Quantification graphs of the effects of OGT overexpression in neurons on intensity, and size of gephyrin or vGAT puncta (Total, non-colocalization). **D.** Quantification of inhibitory synapse number (gephyrin and vGAT puncta) showing no significant changes following OGT deletion. **E.** Quantification of overlapped volume gephyrin and vGAT puncta, showing a significant increase in OGT overexpression neurons. Data are presented as S.E.M (*P < 0.05, **P < 0.01, ***P < 0.001; unpaired Student’s t test).

Quantitative analysis revealed that OGT overexpression reduced several structural parameters of inhibitory synapses (Figure 3B–C). Specifically, vGAT puncta displayed decreased intensity and size in the total pool, whereas vGAT puncta not colocalized with gephyrin were unaffected. The volume of overlap between vGAT and gephyrin was not affected. Similarly, gephyrin puncta exhibited reduced intensity in both the total pool and in non-colocalized fractions. In contrast, the number of neither vGAT nor gephyrin puncta was altered (Figure 3D, E). This indicates that OGT overexpression does not change inhibitory synapse density but specifically alters its structural properties including downregulating inhibitory synapse size.

### OGT knockout increases inhibitory synapse size

Having established that OGT overexpression downregulates inhibitory synapses, we next examined the effects of OGT deletion on inhibitory synapses. Western blot analysis of whole-cell lysates from cultured cortical neurons revealed a significant ∼30% reduction in overall vGAT protein abundance in OGT-KO cultures (Figure 4A-B; P = 0.0035). To measure the synaptic distribution of vGAT and gephyrin, immunocytochemistry (ICC) was used in hippocampal neurons following sparse Cre-mediated OGT deletion. Interestingly, despite the reduction in total vGAT protein observed biochemically, ICC revealed increased vGAT puncta intensity and size (Figure 4D–E). This apparent discrepancy likely reflects a redistribution of available vGAT from the global pool toward synaptic compartments rather than a simple increase in protein abundance. However, vGAT puncta not colocalized with gephyrin were not affected whereas it’s volume of overlap with gephyrin decreased (Figure 4D-E). Notably, total gephyrin intensity and volume were unaffected by OGT deletion. In contrast, OGT deletion was associated with decreased intensity of gephyrin in non-colocalized puncta and increased volume of overlap with vGAT (Figure 4D). These data indicate that removing OGT increases inhibitory synapse size while reducing inhibitory synaptic proteins outside synapses. Similarly, while there was no effect on the number of gephyrin puncta, OGT-KO neurons showed a decrease in non-colocalized vGAT puncta number (Figure. 4G). Together, our results demonstrate that OGT deletion leads to structural remodeling of inhibitory synapses without affecting inhibitory synapse number, with differential effects on presynaptic and postsynaptic compartments.

**Figure 4.**
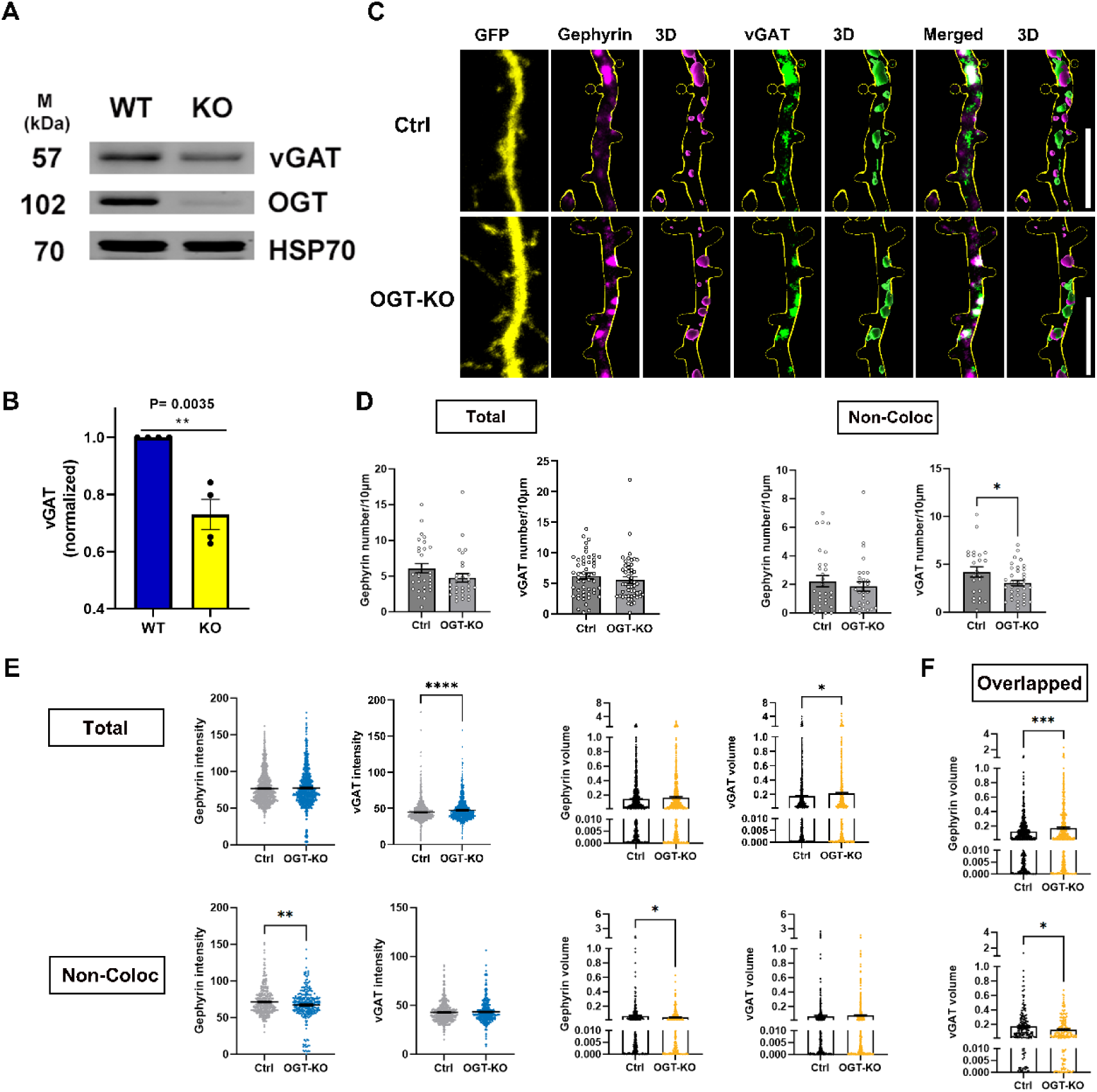
OGT knockout increases inhibitory synapse size. **A.** Representative Western blots showing v-GAT, OGT, and HSP70 protein levels under basal conditions. **B.** Quantification of normalized v-GAT expression showing significant reduction in OGT-KO compared to WT neurons under basal conditions (P = 0.0035; N=4). **C.** Immunocytochemistry of GFP expressing hippocampal neurons stained with antibodies against GFP (green) and OGT (purple) at DIV16. Scale bars 20 μm. **D.** Representative images of cultured hippocampal neurons transfected at DIV7 with expressing GFP alone (Control) or co-expressing GFP with Cre (OGT-KO). Neurons were analyzed by triple immunofluorescence labeling for GFP (green), gephyrin (blue), and vGAT (red). Scale bar, 5 μm (applies to all images). **E.** Quantification graphs of the effects of OGT-KO in neurons on intensity, and size of gephyrin or vGAT puncta (Total, Non-colocalization). Data are presented as S.E.M (*P < 0.05, **P < 0.01, ***P < 0.001; unpaired Student’s t test). **F.** Quantification of inhibitory synapse number (gephyrin and vGAT puncta) showing no significant changes following OGT deletion

### OGT knockout alters inhibitory synaptic transmission kinetics and GABA_A_ receptor subunit composition

To determine whether OGT deletion affects inhibitory synaptic function, we first performed whole-cell recordings of spontaneous inhibitory postsynaptic currents (sIPSCs) in cultured cortical neurons. Analysis revealed that OGT-KO neurons exhibited significantly faster sIPSC decay kinetics compared with WT controls (Figure 5A–C; P = 0.0148), while sIPSC amplitude was not significantly altered (Figure 5B). Faster decay kinetics are consistent with reduced contribution of β3-containing GABA_A_ receptor (Figure 5C), which normally prolong inhibitory current duration[38–40].

**Figure 5.**
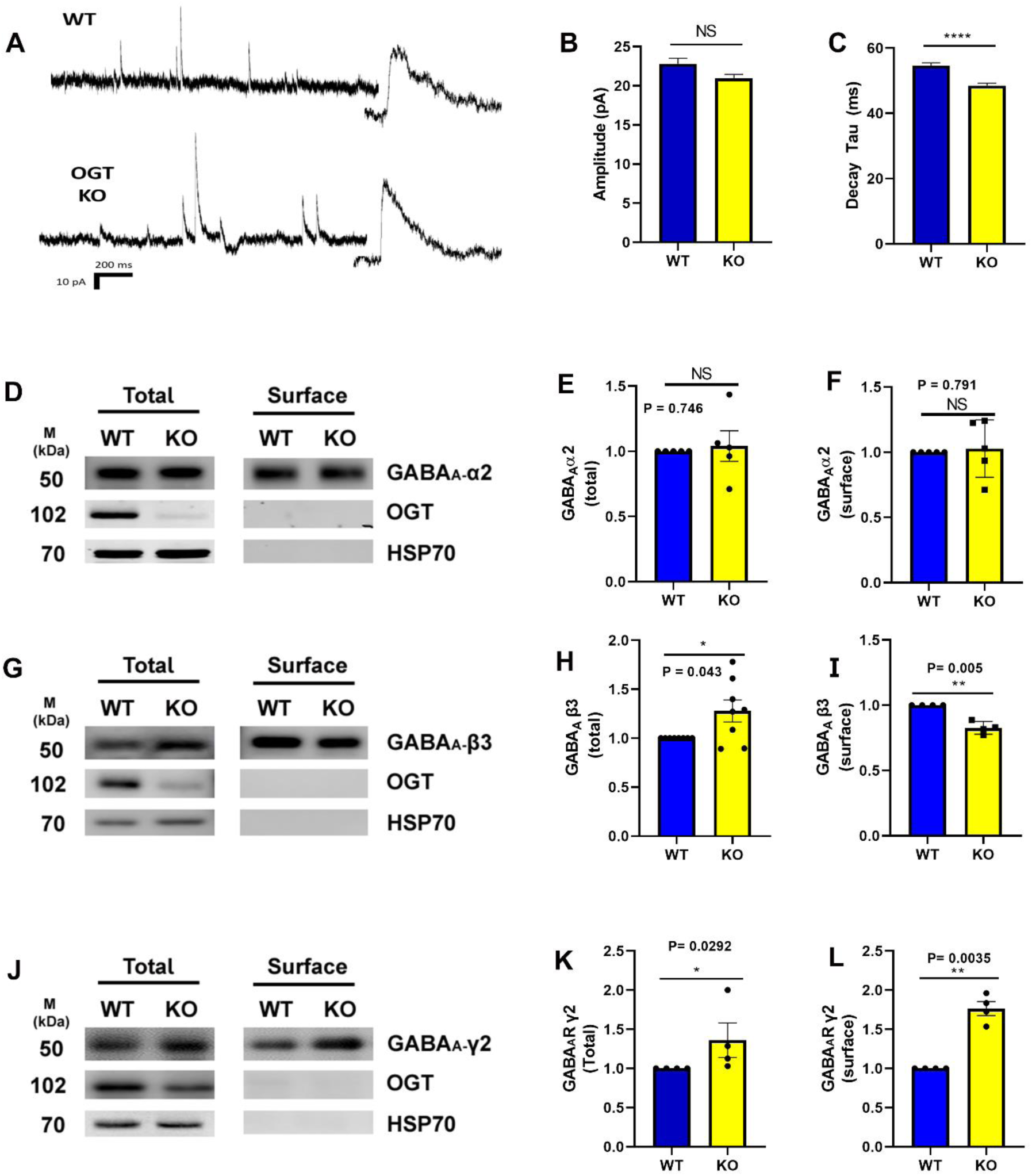
OGT knockout alters inhibitory synaptic transmission kinetics and GABA_A_ receptor subunit composition. **A.** Representative sIPSC traces in WT and OGT-KO(KO) cultured cortical neurons. **B.** sIPSC amplitude in comparing WT vs OGT-KO cultured cortical neurons (NS). **C.** sIPSC decay tau in comparing WT vs OGT-KO cultured cortical neurons (P = 0.0148; Mann-Whitney test). **D.** Representative Western blot analysis of total and surface GABA_A_ α2 subunit expression in wild-type (WT) and OGT knockout (KO) neurons, with OGT and HSP70 included as loading controls. **E.** Quantification of total GABA_A_ α2 subunit protein levels in WT and OGT-KO neurons. **F.** Quantification of surface GABA_A_ α2 subunit protein levels in WT and OGT-KO neurons**. G.** Representative Western blot analysis of total and surface GABA_A_ β3 subunit expression in WT and KO neurons, with OGT and HSP70 included as loading controls. **H.** Quantification of total GABA_A_ β3 subunit protein levels in WT and OGT-KO neurons. **I.** Quantification of surface GABA_A_ β3 subunit protein levels in WT and OGT-KO neurons. **J.** Representative Western blot analysis of total and surface GABA_A_ γ2 subunit expression in WT and KO neurons, with OGT and HSP70 included as loading controls. **K.** Quantification of total GABA_A_ γ2 subunit protein levels in WT and OGT-KO neurons. **L.** Quantification of surface GABA_A_ γ2 subunit protein levels in WT and KO neurons. All statistical analyses were performed using the Mann-Whitney test.

Building on this functional change, we next investigated whether OGT deletion alters expression of specific GABA_A_ receptor subunits, the principal mediators of fast inhibitory signaling [41]. We focused on three subunits with well-established functional relevance: α2, which contributes to synaptic targeting and confers relatively fast decay kinetics[42]; β3, which regulates current kinetics and surface trafficking[41]; and γ2, which is essential for receptor clustering at inhibitory synapses and present in ∼90% of synaptic GABA_A_ receptors[43, 44].

Western blot analysis of total and biotinylated surface protein fractions revealed distinct, subunit-specific patterns of regulation by OGT (Figure 5D-L). The α2 subunit showed no significant changes in either total or surface expression levels following OGT deletion, indicating that this subunit is not directly regulated by OGT (Figure 5D-F). However, the β3 subunit displayed a complex regulatory pattern, with significantly increased total protein levels (P = 0.043) but decreased surface expression in OGT-KO neurons (P = 0.005) compared to controls (Figure 5G-I). This opposing effect suggests that while OGT normally limits total GABA_A_ β3 synthesis, it simultaneously facilitates the trafficking of β3-containing receptors to the plasma membrane. In contrast, the γ2 subunit showed robust upregulation in both total and surface fractions of OGT-KO neurons (Figure 5J-L). Total γ2 expression was significantly elevated (P = 0.029). Surface γ2 levels demonstrated an even more pronounced increase (P = 0.0035). These findings indicate that OGT has a wide-spread and diverse effect on GABA_A_ receptor regulation.

### OGT knockout affects GABA_A_ synaptic organization

Our biochemical data indicate that OGT regulates the synaptic levels of GABA_A_ receptors. Testing this hypothesis, we performed ICC using surface staining of γ2 hippocampal neurons following sparse OGT deletion (Figure 6A). We co-stained for vGAT to distinguish between synaptic and extra synaptic receptors. Imaris-based analysis of GFP-labeled dendrites showed that OGT-KO neurons did not exhibit any change in the number of γ2 receptor puncta (Figure 6B). However, OGT-KO significantly increased the total γ2 puncta intensity puncta intensity. Similarly, there was also an increase in the volume of γ2 that overlapped with vGAT. γ2 puncta volume for both total and non-colocalized with vGAT also showed an increased trend in OGT-KO neurons compared to wildtype, although they did not reach statistical significance (Figure 6C). In addition, there was an increase in the volume of γ2 that overlapped with vGAT. Consistent with these postsynaptic γ2 changes, OGT-KO neurons displayed a significant increase in the overlapped volume between vGAT and γ2 (Figure 6D).

**Figure 6.**
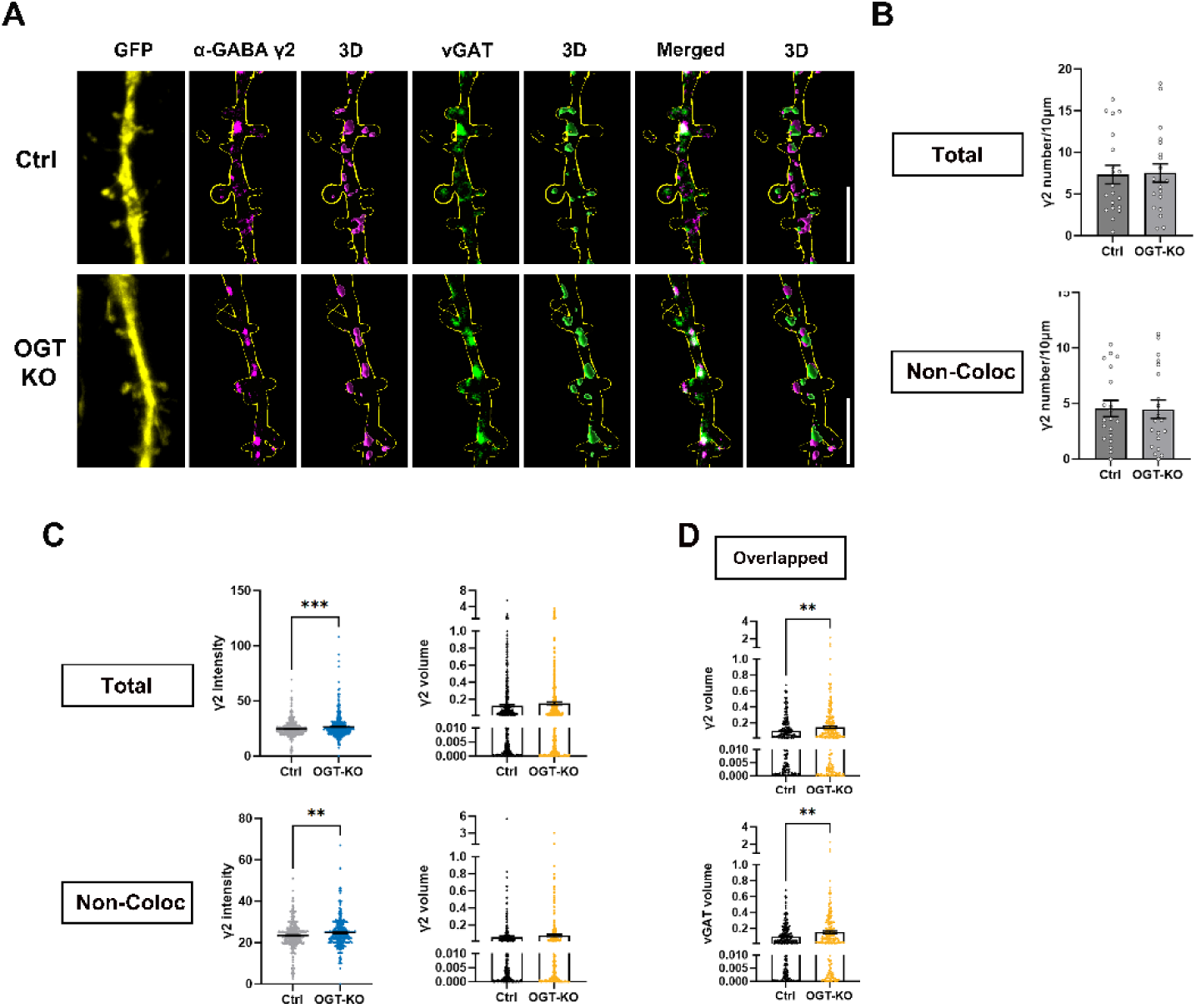
OGT knockout affects synaptic organization. **A.** Representative images of cultured hippocampal neurons transfected at DIV7 with GFP alone (Control) or GFP + Cre (OGT-KO). Neurons were analyzed by triple immunofluorescence labeling for GFP (yellow), α-GABA_A_ γ2 (magenta), and vGAT (green). Shown are single-plane and 3D-rendered views of γ2 and vGAT puncta along dendritic segments, with merged images illustrating their spatial relationship. Scale bar, 5 μm (applies to all images). **B**. Quantification of γ2 and vGAT puncta number per 10 μm dendrite in control and OGT-KO neurons, shown for total and non-colocalized populations. No significant differences were observed**. C.** Quantification of γ2 puncta intensity and volume (total and non-colocalized with vGAT) showed that OGT-KO neurons exhibited significant increases in γ2 intensity compared with controls. **D.** Quantification of overlapped γ2–vGAT puncta volume, showing significantly increased overlap in OGT-KO neurons. Data are presented as S.E.M. *P < 0.05, **P < 0.01, ***P < 0.001; unpaired Student’s t test.

In concurrence with the measurements of morphological synapse number and biochemical GABA_A_ receptor expression, these observations indicate that OGT regulates both the synaptic and extra synaptic levels of γ2, without affecting inhibitory synapse number. Together, our results demonstrate that OGT regulates the structure and function of inhibitory synapses without affecting their number.

## Discussion

Our study shows that OGT regulates inhibitory synapse structure and function. We demonstrate that OGT deletion reduces basal c-Fos expression yet strongly enhances responsiveness to disinhibition, pointing to an important role for OGT in regulating inhibitory control (Figure 1). Manipulation of OGT produced bidirectional structural changes (Figures 3–4), altered inhibitory current kinetics and GABA_A_ receptor subunit composition without affecting synapse number (Figures 5 and 6). These effects were cell-autonomous and mediated at least in part through postsynaptic mechanisms. Indeed, OGT localized to postsynaptic inhibitory terminals (Figure 2). Together, our results identify OGT as a synaptic regulator of inhibitory synapse structure and receptor distribution. By linking metabolic signaling to inhibitory synapse remodeling, these findings show how OGT helps coordinate excitatory and inhibitory balance, providing a mechanism through which internal metabolic state can shape circuit function.

Little is known about what mechanisms regulate inhibitory synapses. Prior work has shown that increasing O-GlcNAc in part by stimulating metabolic flux through the HBP suppressed hippocampal GABA currents [26]. However, whether and how O-GlcNAc may regulate inhibitory synapses has been unclear.

Manipulating OGT expression altered inhibitory synapse structure in a bidirectional manner. We measured effects on synapse structure by imaging the presynaptic protein vGAT and the postsynaptic protein gephyrin. Overexpression reduced both vGAT and gephyrin in dendrites (Figure 3C–E). In contrast, knockout increased vGAT puncta size and intensity (Figure 4E–F). These results are consistent without imaging of surface γ2 receptors: deleting OGT increases γ2 puncta intensity and overlap with vGAT without affecting the γ2 puncta number along the dendrite. However, there was no change in puncta number in any condition except for the number of non-colocalized vGAT puncta (Figure 3D, 4G). The opposing outcomes of overexpression and knockout converge on OGT as a regulator of inhibitory synapse structure, rather than synapse number, aligning with nanoscale models of inhibitory remodeling [45]. The increase in gephyrin overlap and decrease in vGAT overlap suggest also that inhibitory synapse organization and alignment depend on OGT. While our data show that OGT-dependent remodeling of inhibitory synapses that involves both pre- and postsynaptic elements, our microscopy data were obtained by postsynaptic OGT manipulation in sparse neurons. This distinction indicates that OGT-dependent changes in the postsynaptic compartment may influence presynaptic protein distribution, possibly through retrograde signaling, a mechanism observed in other forms of synaptic plasticity [8, 46]. Together, the data favor a model where OGT restrains inhibitory synapse structure.

In addition to remodeling inhibitory synapse structure, OGT regulates the distribution of inhibitory synaptic proteins between synaptic and extrasynaptic compartments. OGT deletion decreased gephyrin that did not colocalize with vGAT but increased the volume of gephyrin that overlapped with vGAT. This indicates that deleting OGT redistributed gephyrin from extra- to intrasynaptic compartments. Similarly, our data are consistent with a redistribution of vGAT from the global pool toward functional synapses in OGT KOs: biochemical analysis showed reduced total vGAT protein (Figure 4A–B) and imaging fewer vGAT puncta not colocalizing with gephyrin (Figure 4D) whereas dendritic vGAT puncta intensity and volume increased as well as its overlap with γ2 (Figure 6C). γ2-containing receptors increased in both extra- and synaptic pools (Figure 6C–D), with greater alignment with vGAT (Figure 6E–F). These data suggest that deleting OGT promotes a redistribution of inhibitory proteins from extrasynaptic sites to synapses.

Consistent with a synaptic effect, our electrophysiological recordings showed faster sIPSC decay kinetics in OGT-KO neurons compared with WT (Figure 5A, C). This indicates that OGT regulates GABA_A_ receptor distribution, in particular reduced β3 contribution to synaptic assemblies [38–40]. Biochemical analysis supported this finding, showing decreased β3 surface expression (Figure 5G–I) together with increased γ2 expression at both total and surface levels (Figure 5J–L). Imaging data in OGT-KO neurons further substantiated the biochemical findings, showing increased γ2 intensity at synaptic and extrasynaptic sites with enhanced alignment to vGAT (Figure 6C–F). These subunit-specific changes in receptor composition provide a mechanistic basis for the altered decay kinetics. The unchanged α2 levels (Figure 5D–F) suggest that OGT does not directly regulate α2 but selectively regulates β3 and γ2 subunits, each of which is known to control inhibitory current kinetics and clustering at synapses [9–12]. As further evidence that OGT regulates inhibitory neurotransmission, bicuculline, a GABA_A_ receptor antagonist elicited an exaggerated c-Fos response in OGT-KO neurons (Figure 1E–F). This indicates that deleting OGT increases inhibitory input. The finding is in concordance with decreased inhibitory neurotransmission in the hippocampus upon metabolic stimulation of O-GlcNAc [26]. As our data shows OGT localizes at inhibitory synapses (Figure 2) and regulates inhibitory synapse remodeling, it raises the possibility that OGT regulates inhibitory neurotransmission by direct modification of synaptic proteins such as gephyrin, γ2 or β3. Taken together, the electrophysiological, imaging and biochemical results demonstrate a direct link between OGT-dependent GABA_A_ receptor composition and inhibitory current properties.

These results on inhibitory synapse structure and function change the model of how neural circuits can integrate metabolic fluctuations. Work from us and others have shown repeatedly that OGT in response to nutritional challenges affects excitatory neurotransmission through the AMPA receptor [14–17]. Preliminary results from the hippocampus indicated that metabolic stimulation of O-GlcNAc may affect not only excitatory but also inhibitory neurotransmission [26]. Neuronal O-GlcNAc cycling is regulated by physiological fluctuations in glucose, metabolic hormones and behavior that affects metabolic state such as feeding [47, 48]. Our data on inhibitory synapse remodeling identifies a molecular mechanism by which OGT can balance excitatory with inhibitory input. Its effect on inhibitory synapses differs in part from how it affects excitatory synapes. While OGT promotes dendritic spine and excitatory synapse number[16], OGT was observed to restrain inhibitory synapse organization without affecting the number of inhibitory synapses. In contrast, the present findings indicate that OGT in both excitatory and inhibitory synapses regulates a neurotransmitter receptor switch [16]. Neurexin and a few other adhesion molecules are known to affect both excitatory and inhibitory synapses [49]. However, our results reveal a mechanism that can tune neural circuit function to the metabolic state of the body by coordinating excitatory with inhibitory input.

Coordination of excitatory and inhibitory input is a fundamental property of neuronal circuits. Its dysregulation is associated with a number of brain disorders, including autism, schizophrenia and intellectual disability[7, 30, 50]. Indeed, OGT has been coupled in humans to brain development, learning and memory and adaptive behavior[51]. Our findings establish OGT as a regulator of inhibitory synapse organization and suggest that it functions as a shared postsynaptic modulator across excitatory and inhibitory synapses. This work provides a foundation for mechanistic dissection of how OGT balances neural circuit function in health and disease.

## Materials and Methods

### Animals and OGT-floxed Mice

Mice carrying loxP sites flanking exon 4 of the Ogt gene (OGT^Fl/^Fl) were maintained on a C57BL/6J background. Breeding pairs were genotyped by PCR as described previously[52]. Animals were housed under a normal 12 h light–dark cycle with ad libitum water and food. The ambient temperature was 25 °C and humidity was 50%. Both male and female mice were used in the study. We have complied with all relevant ethical regulations for animal use. All experimental procedures were approved and conducted in accordance with the regulations of the Local Animal Ethics Committee at Umeå University.

### Lentivirus Production

To generate lentiviral constructs for OGT knockout in cultured neurons, pseudotyped vesicular stomatitis virus G (VSV-G) lentiviruses expressing either GFP alone (wild-type control) or GFP together with Cre recombinase (OGT knockout) were produced following standard protocols. HEK293T cells were co-transfected with the transfer vector (FUGW or FUGW-Cre), the packaging plasmid (pCMVΔR8.9), and the envelope plasmid (pMD2.G-VSV-G) using Lipofectamine 2000 (Invitrogen, #12313563) according to the manufacturer’s instructions. Viral supernatants were collected 48–64 h post-transfection, filtered through 0.45 µm filters, and concentrated by ultracentrifugation. Viral aliquots were stored at −80 °C until use.

### Primary Neuronal Culture

Primary cortical and hippocampal neurons were prepared from embryonic day 16.5 (E16.5) Ogt^fl/fl^ or wild-type littermate mice, as described previously[15, 16, 52]. Forebrains were dissected in ice-cold Hank’s balanced salt solution (HBSS) and enzymatically dissociated with papain (20 U/mL; Worthington) for 20 min at 37 °C. Cells were gently triturated in NM5 (Neurobasal medium (Gibco, #11570556) supplemented with 2% (v/v) B27 (Invitrogen, #15360284), 2 mM GlutaMAX (Gibco, #35050-038), 5%(v/v) fetal bovine serum (FBS)(Cytiva, #10309433), and 100 U/mL penicillin/streptomycin(Pen-Strep)(Gibco, #15140-122).

Dissociated cortical neurons were plated on poly-D-lysine– or poly-L-lysine–coated 35 mm dishes or coverslips at a density of 1 × 10⁶ cells per well. After 2 h, plating medium (NM5) was replaced with fresh NM5 medium. Cultures were maintained at 37 °C in a humidified incubator (5% CO₂). At DIV3–5, glial proliferation was suppressed by adding 5 µM 5-fluoro-2′-deoxyuridine (FDU). From this point on, half of the medium was replaced every 3–4 days with glia-conditioned Neurobasal medium containing 1% FBS and 2% B27. For genetic manipulation, cortical neurons were transduced at DIV2 with lentiviral vectors encoding Cre-GFP (for OGT knockout) or GFP alone (control). Cortical neurons were used between DIV14–18.

Dissociated hippocampal neurons were plated on poly-L-lysine–coated coverslips at a density of 3 × 10⁵ cells per well. After 2 h, plating medium (NM5) was replaced with NM0 medium (Neurobasal + GlutaMAX™ + Penstrep + 2% B27). Cultures were maintained at 37 °C (5% CO₂), with no further media changes until the time of transfection. Hippocampal neurons were transfected at DIV7 using the appropriate DNA constructs. These neurons were also used for imaging experiments 48-hours post-transfection.

### Transfection

At DIV7, primary hippocampal neurons cultured in 12-well plates were transfected with 1.5 µg of plasmid DNA each well. Lipofectamine (2 µl per well; Thermo Fisher Scientific, 10696343) and DNA were each diluted to 50 µl per well in plain Neurobasal medium, incubated separately for 5 minutes, then combined at a 1:1 ratio and incubated for an additional 20 minutes at 37 °C. The transfection complexes (100 µl per well) were applied to the neurons for 45 minutes, then all wells were rinsed with plain Neurobasal medium and returned to conditioned medium.

### Immunochemistry

Cultured neurons were fixed with 4% paraformaldehyde and 4% sucrose in ice-cold PBS for 15 minutes at 4 °C. Cells were then permeabilized with 0.25% Triton X-100 for 10 minutes at 4 °C (permeabilization was omitted for surface staining) and blocked for 1 hour at room temperature in blocking buffer (10% Normal goat serum, Vector Laboratories, S-1000-20). Subsequently, neurons were incubated with primary antibodies for 2 hours at room temperature. After washing, secondary antibodies were applied for 1 hour at room temperature. Following three washes (10 minutes each) in PBS, samples were mounted using fluorescence mounting medium (Fluoromount-G, Thermo Fisher Scientific, 15586276) and stored at 4 °C.

Antibodies for immunocytochemistry: anti-GFP (Aves labs, GFP-1020, 1:1000), anti-OGT (Proteintech, 11576-2-AP, 1:1000), anti-Gephyrin (Synaptic Systems, 147011, 1:1000), anti-vGAT (Synaptic Systems, 131005, 1:1000), anti-α-GABAA γ2 (Synaptic Systems, 224003, 1:1000), anti-Chicken secondary antibody(Azure Biosystems, AC2209, 1:2000), anti-Guinea Pig secondary antibody(Thermo Fisher Scientific, A21450, 1:2000), anti-Mouse secondary antibody (Thermo Fisher Scientific, A21422, 1:2000), anti-Rabbit secondary antibody(Thermo Fisher Scientific, A21428, 1:2000).

### Image acquisition and analysis

Confocal images of neuronal cultures and brain sections were acquired using a Leica SP8 FALCON confocal microscope. Z-stack images spanning approximately 50 µm in thickness were collected at 0.5-µm intervals. Image analysis was performed using Imaris 10.2.1 (Bitplane). The number, volume, and intensity of protein marker–positive puncta were quantified. Gephyrin, vGAT and α-GABAA γ2 within the GFP-positive neurons were three-dimensionally reconstructed and automatically analyzed using Imaris software.

Filament and surface reconstruction in Imaris quantified three primary parameters: intensity (fluorescence signal strength, reflecting relative protein abundance), volume (3D puncta size), and puncta number. ‘Total puncta’ refer to the entire set of immunoreactive structures within the analyzed dendritic segments, including both synaptic and extrasynaptic clusters. Puncta were additionally categorized as non-colocalized (single-marker puncta lacking an apposed partner, representing isolated pre- or postsynaptic or extrasynaptic sites) or overlapped puncta, for which Imaris calculated the shared 3D volume between vGAT and gephyrin, representing structural alignment between presynaptic boutons and postsynaptic scaffolds.

To ensure consistent presentation, moderate linear adjustments were applied uniformly across entire images (not to specific regions). All images were acquired under identical microscope settings and processed in the same manner using Leica X Office software.

### Bicuculline Treatment

To stimulate neuronal network activity by blocking inhibitory neurotransmission, cortical cultures at DIV14–18 were treated with bicuculline (40 μM; Sigma-Aldrich, #14340) for 4 h at 37 °C to pharmacologically inhibit GABA_A_ receptors. Following treatment, cultures were processed immediately for subsequent experimental procedures. Control cultures received vehicle treatment under identical conditions.

### Surface Biotinylation and Pulldown

At 14–18 days in vitro (DIV14–18), cultured neurons were washed three times with ice-cold artificial cerebrospinal fluid (aCSF; 143 mM NaCl, 5 mM KCl, 2 mM CaCl₂, 1 mM MgCl₂, 10 mM HEPES, pH 7.4, and 10 mM D-glucose). Surface proteins were labeled by incubating cells on ice for 20 min with 1 mg/mL Sulfo-NHS-SS-Biotin (Fisher Scientific, 15332627) prepared in aCSF. Excess biotin was quenched twice with 50 mM glycine in aCSF for 5 min each. Cells were then lysed in neural lysis buffer containing 50 mM Tris–Cl (pH 7.4), 150 mM NaCl, 1% (v/v) Triton X-100, and 0.1% (w/v) SDS, supplemented with protease, phosphatase, and O-GlcNAcase inhibitors. Lysates were clarified by centrifugation at 14,000 × g for 10 min at 4°C, and the supernatants were incubated overnight at 4 °C with NeutrAvidin agarose beads (Thermo Fisher Scientific) to isolate biotinylated surface proteins. Beads were washed three times with neural lysis buffer and eluted with 2× Laemmli sample buffer containing 50 mM DTT by heating at 45 °C for 10 min, as recommended by the antibody manufacturer to prevent protein aggregation. Eluted samples were analyzed by SDS–PAGE and Western blotting.

### Western Blot

Homogenized brain tissue was lysed in neural lysis buffer containing 50 mM Tris–Cl (pH 7.4), 150 mM NaCl, 1% (v/v) Triton X-100, and 0.1% (w/v) SDS, supplemented with protease, phosphatase, and O-GlcNAcase inhibitors. Lysates were cleared by centrifugation at 13,000 rpm for 15 min at 4 °C, and the supernatants were collected for protein analysis. Protein samples were mixed with loading buffer and denatured by boiling for 10 min at 90 °C. For blots detecting GABA_A_ receptor subunits, samples were denatured for 15 min at 45 °C, as recommended by the antibody manufacturer to prevent aggregation.

Western blotting was performed using standard procedures. Equal amounts of protein were separated by SDS–PAGE and transferred onto polyvinylidene fluoride (PVDF) membranes. Membranes were blocked in 5% non-fat milk prepared in TBS-T and incubated overnight at 4°C with primary antibodies against OGT (Proteintech, 11576-2-AP; 1:2000), c-Fos (Santa Cruz, sc-52; 1:1000), HSP70 (Proteintech, 10995-1-AP; 1:10,000), GABA_A_ receptor γ2 (Synaptic Systems, 224 003; 1:1000), α2 (Synaptic Systems, 224 102; 1:1000), β3 (NeuroMab, N87/25; 5µg/blot), or vGAT (Synaptic Systems, 131 013; 1:1000). After washing, membranes were incubated with HRP-conjugated secondary antibodies (anti-rabbit, Thermo Fisher Scientific, 31460; or anti-mouse, Thermo Fisher Scientific, 31430) for 1 h at room temperature. Protein bands were visualized using enhanced chemiluminescence detection reagent (Thermo Fisher Scientific, 34580) and imaged with a Sapphire Biomolecular Imager (Azure Biosystems, IS4000). Band intensities were quantified using Image J software and normalized to HSP70 as a loading control for homogenate samples.

### Electrophysiology

Perforated whole-cell patch-clamp recordings were performed on primary cortical neurons at DIV8-11 from wild-type (WT) and OGT knockout (OGT-KO) cultures. Neurons were visually identified using an epifluorescence microscope (Zeiss Axio Examiner) equipped with a monochromator and TILLVisIon software (T.I.L.L. Photonics GmbH, Munich, Germany). GFP-positive cells were selected for recordings to ensure viral transduction and genotype identity.

Patch pipettes were pulled from borosilicate glass capillaries (Harvard Apparatus, UK) using a Flaming/Brown micropipette puller (Model P-97, Sutter Instrument Co., USA) to a resistance of 3–5 MΩ when filled with internal solution. The pipette solution contained (in mM): 140 CsAc, 3 NaAc, 1.2 MgCl_2_, 1 EGTA, 10 HEPES, 10 sucrose (pH adjusted to 7.2 with CsOH) and supplemented with 400 µM amphotericin B (Sigma-Aldrich) dissolved in DMSO.

The extracellular recording solution contained (in mM): 119 NaCl, 5 KCl, 2.5 CaCl₂, 1.5 MgCl₂, 20 HEPES, 25 D-glucose and 5 sucrose (pH 7.4, adjusted with NaOH). Recordings were performed at room temperature using a MultiClamp 700B amplifier (Molecular Devices, USA). Neurons were voltage-clamped at 0 mV, and spontaneous inhibitory postsynaptic currents (sIPSCs) were recorded in the absence of synaptic blockers. Signals were low-pass filtered at 2 kHz, digitized at 10 kHz, and acquired with Clampex software (Molecular Devices).

Data were analyzed using Clampfit 10.7 (Molecular Devices) and Mini Analysis (Synaptosoft). The decay kinetics of sIPSCs were fitted with a single exponential function to determine the decay time constant (τ).

### Statistical Analysis

Data represent mean ± SEM from at least three independent cultures. Data are presented as mean ± SEM. Statistical analyses were performed using GraphPad Prism 10. For normally distributed data, unpaired two-tailed Student’s t-tests were used. For non-normally distributed data, Mann–Whitney U tests were applied. P < 0.05 was considered statistically significant. Exact n values and statistical tests used for each experiment are reported in the figure legends.

## Acknowledgments

We thank the staff at the Umeå Centre for Comparative Biology (UCCB) for excellent animal care and technical assistance. We acknowledge the Biochemical Imaging Center at Umeå University and the National Microscopy Infrastructure, NMI (VR-RFI 2023-00163) for providing assistance in microscopy

## Funding and additional information

This work was supported by the Knut and Alice Wallenberg Foundation (OL), the Swedish Scientific Council (grant no. 2022-01024, OL), Umeå University, the Swedish Brain Foundation, Region Västerbotten, the Kempe Foundations, Märta Lundqvist’s Foundation, Fredrik och Ingrid Thuring’s Foundation, and the Sigurd och Elsa Golje Memory Foundation.

